# Changes in barnacle recruitment and recruit body size with intertidal elevation on wave-exposed rocky shores in Nova Scotia

**DOI:** 10.1101/766618

**Authors:** Ricardo A. Scrosati

## Abstract

Barnacle recruitment is often studied in rocky intertidal habitats due to the relevant role that barnacles can play in intertidal communities. In 2014, barnacle (*Semibalanus balanoides*) recruitment was measured at high elevations in wave-exposed intertidal habitats on the NW Atlantic coast in Nova Scotia, Canada. Values were considerably lower than previously reported for middle elevations in wave-exposed intertidal habitats on the NE Atlantic and NE Pacific coasts. To determine if such differences in recruitment may have resulted from elevation influences, I did a field experiment in 2019 in wave-exposed intertidal habitats in Nova Scotia to test the hypothesis that recruitment is higher at middle than at high elevations, based on known environmental differences between both elevation zones. Based on data from three locations spanning 158 km of the Nova Scotia coast, barnacle recruitment was, on average, nearly 200 % higher (and recruits were larger) at middle than at high elevations. However, even with this increase, barnacle recruitment on this NW Atlantic coast is still lower than for comparable habitats on the NE Atlantic and NE Pacific coasts, and also lower than previously reported for wave-exposed locations farther south on the NW Atlantic coast, in Maine, USA. Therefore, barnacle recruitment in wave-exposed intertidal environments in Nova Scotia appears to be only moderate relative to other shores. This difference in the supply of barnacle recruits might influence the intensity of interspecific interactions involving barnacles.

## Introduction

Barnacles are common organisms in rocky intertidal habitats worldwide (Anderson 1994). For these organisms, recruitment refers to the appearance of new individuals in intertidal habitats as a result of the metamorphosis of pelagic larvae that have settled on the intertidal substrate. Besides ensuring the long-term persistence of barnacle populations, barnacle recruitment can also positively influence other intertidal species, such as mussels by facilitating their own recruitment (Menge 1976) and predatory snails by providing them with an accessible food source (Noda 2004). In studies on the influence of coastal waters on intertidal communities, barnacle recruitment has often been measured due to its sensitivity to nearshore pelagic conditions and its influence on intertidal community structure (Cole et al. 2011, Jenkins et al. 2000, Mazzuco et al. 2018, Menge and Menge 2013, Navarrete et al. 2008, Shanks et al. 2017). Studies of that kind have recently began on the NW Atlantic coast in Canada (Scrosati and Ellrich 2018).

Specifically, barnacle recruitment (recruit density) was measured in 2014 in wave-exposed rocky intertidal habitats on the Atlantic coast of Nova Scotia (Scrosati and Ellrich 2018). Although that study measured barnacle recruitment at various locations, values were nonetheless considerably lower than those reported for many wave-exposed habitats on the NE Atlantic (Jenkins et al. 2000) and NE Pacific (Navarrete et al. 2008) coasts. Such a disparity between coasts tentatively suggests that barnacles might play a more limited ecological role in Nova Scotia. Investigating this possibility seems, in principle, appealing to learn more about barnacle influences in intertidal communities. However, the observed disparity in recruitment between these coasts might simply have resulted from basic sampling differences. For instance, the Nova Scotia study measured barnacle recruitment at high intertidal elevations (Scrosati and Ellrich 2018), while the studies done on the other coasts measured recruitment at middle (Jenkins et al. 2000) and middle-to-low (Navarrete et al. 2008) intertidal elevations. Since middle elevations spend more time underwater than high elevations because of tide dynamics, middle elevations experience lower desiccation and thermal extremes during low tides and offer a longer time for larval settlement during high tides than high elevations (Menge and Branch 2001, Raffaelli and Hawkins 1999). Thus, barnacle recruitment could be higher at middle than at high intertidal elevations in Nova Scotia. To test this hypothesis, I conducted a field experiment in 2019 on the Atlantic coast of Nova Scotia.

Besides recruit density, recruit size is another trait of potential ecological relevance, as larger body sizes of prey represent more food for predators (Carroll and Wethey 1990, Dunkin and Hughes 1984). The lower desiccation and thermal extremes during low tides and the longer availability of pelagic food during high tides at middle elevations than at high elevations (Menge and Branch 2001, Raffaelli and Hawkins 1999) predict larger barnacle recruits at middle elevations (given the same time for growth; Wethey 1983). I tested this hypothesis also through the experiment done in 2019 on the Atlantic coast of Nova Scotia.

## Materials and methods

I performed the experiment between April and July 2019 in rocky intertidal habitats on the open Atlantic coast of Nova Scotia, specifically at three locations spanning 158 km along the coast: Duck Reef (44° 29’ 29” N, 63° 31’ 37” W), Western Head (43° 59’ 23” N, 64° 39’ 39” W), and West Point (43° 39’ 12” N, 65° 07’ 51” W), from north to south (Fig. 1). The surveyed intertidal substrate was stable bedrock in all cases. These are wave-exposed intertidal locations, as they face the open Atlantic Ocean without any obstructions. Daily maximum water velocity (an indication of wave exposure) measured with dynamometers in wave-exposed intertidal habitats on this coast ranges between 6–12 m s^−1^ (Ellrich and Scrosati 2017, Hunt and Scheibling 2001, Scrosati and Heaven 2007).

**Figure 1.**
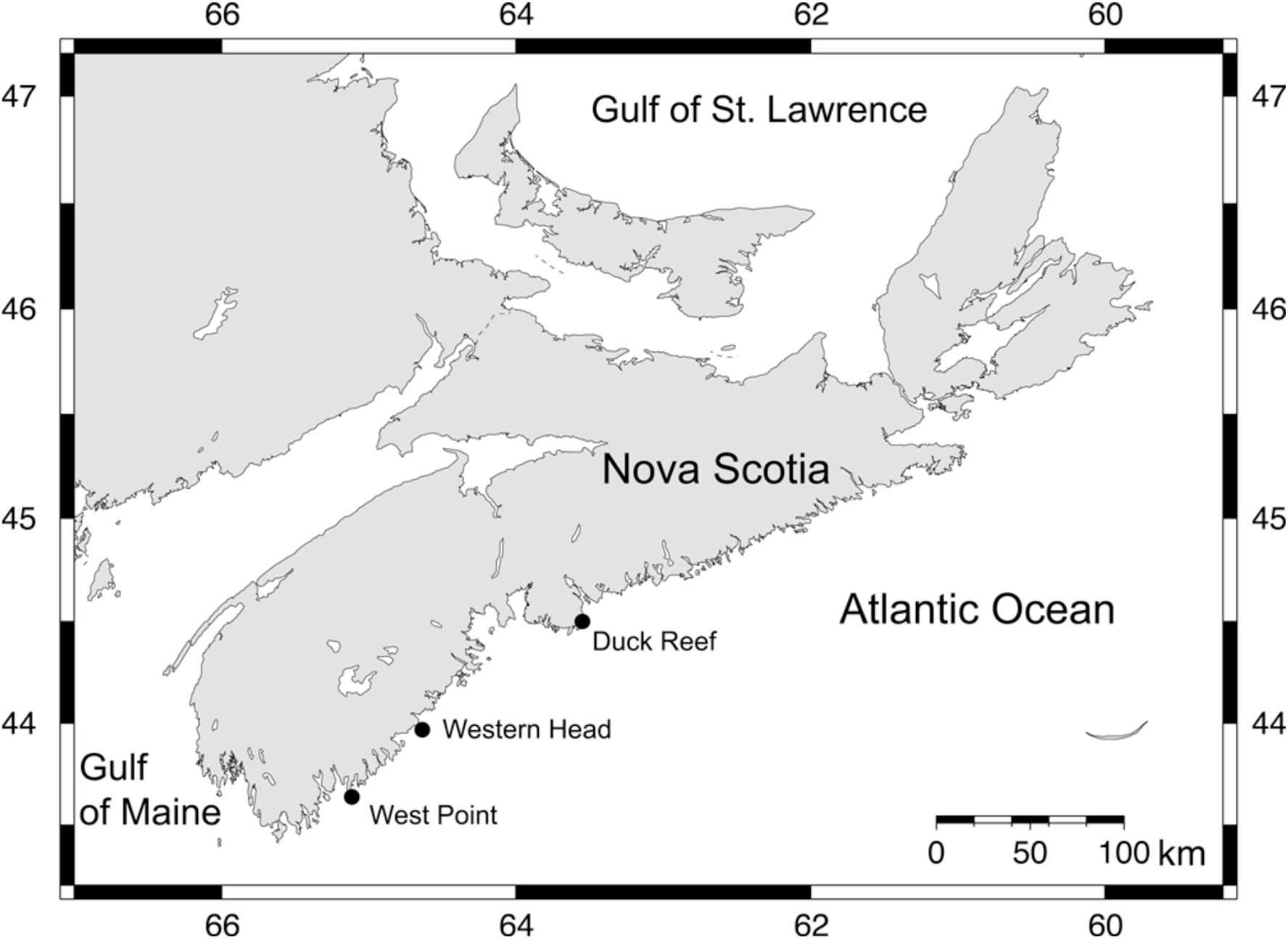
Map indicating the position of the three wave-exposed intertidal locations studied on the Atlantic coast of Nova Scotia, Canada.

Along the studied coastal range, tidal amplitude increases southwards from 2.1 m to 2.4 m (Tide-Forecast 2019). In addition, even though the three studied locations face the open ocean directly, small differences in wave exposure between them may have existed (influenced by, for example, local topography). Thus, to ensure that all barnacle recruitment data intended to represent either “high intertidal” or “mid-intertidal” conditions were taken at a comparable elevation (in terms of exposure to aerial conditions during low tides) across the three locations, I used a sampling procedure based on a biological indicator (Scrosati and Heaven 2007). At each location, I measured the intertidal range as the vertical distance between chart datum (0 m in elevation) and the highest elevation where perennial sessile organisms (coincidentally, the studied barnacle, *Semibalanus balanoides* (L.)) occurred on the substrate outside of crevices. Then, for each location, I divided the vertical intertidal range in three and measured barnacle recruitment at an elevation just above the bottom boundary of the upper third of the intertidal range (representing “high intertidal” conditions) and at a lower elevation 30–40 cm below (representing “mid-intertidal” conditions). The sampled elevation representing high intertidal conditions is the same as that one sampled in 2014 (Scrosati and Ellrich 2018).

The studied barnacle, *Semibalanus balanoides*, is the only intertidal species of barnacle on this coast. In Atlantic Canada, *S. balanoides* mates in autumn, broods in winter, and releases pelagic larvae in spring (Bouchard and Aiken 2012, Bousfield 1954, Crisp 1968). Larvae settle in intertidal habitats and metamorphose into benthic recruits in May and June, which is thus considered to be the recruitment season (Ellrich et al. 2015). To measure barnacle recruitment and body size unaffected by other sessile species (Beermann et al. 2013), I made clearings (100-cm^2^ quadrats) of the natural rocky substrate at each location in late April, eight at the high intertidal zone (except at Western Head, where six were made) and eight at the middle intertidal zone. I measured barnacle recruit density and size between 2-4 July using digital photographs of the quadrats taken at low tide. I calculated barnacle recruit density as the number of barnacle recruits found in a quadrat divided by quadrat area (1 dm^2^). I calculated recruit size as the basal shell diameter measured along a straight line passing through the middle of the rostrum and the carina (Chan et al. 2006). I measured size for a maximum of 10 recruits per quadrat (all recruits if there were 10 or less in a quadrat and a random selection of 10 if there were more than 10 in a quadrat). I then calculated mean recruit size for each quadrat, using those values as replicates for the data analysis. Finally, I evaluated the interactive effects of intertidal elevation (fixed factor with two levels) and location (random factor with three levels) on barnacle recruit density and size separately for each of these two dependent variables through a two-way analysis of variance (Quinn and Keough 2002).

## Results

On 2 May, a visual inspection of the quadrats revealed that no barnacle recruits were then present on the substrate. On 9 May, another inspection indicated that recruits had already began to appear at both the high and mid-intertidal zone. Further visual inspections indicated that new recruits kept appearing in the quadrats in the following weeks (mostly during May). No new recruits appeared after 24 June. No dead recruits (empty shells attached to the substrate) were found during these inspections or in early July when the quadrats were photographed. Therefore, all of the recruits that progressively appeared in the quadrats during the recruitment season seem to have contributed to the values of barnacle recruit density measured in early July.

Based on the data collected on 2-4 July, barnacle recruit density in 2019 was significantly higher at the mid-intertidal zone than at the high intertidal zone (*F*_1, 2_ = 44.91, *P* = 0.022), 197 % higher on average (Fig. 2). This outcome was consistent across locations, as the interaction term between elevation zone and location was not significant (*F*_2, 40_ = 0.21, *P* = 0.808). Barnacle recruit density was statistically similar among locations (*F*_2, 40_ = 3.12, *P* = 0.055).

**Figure 2.**
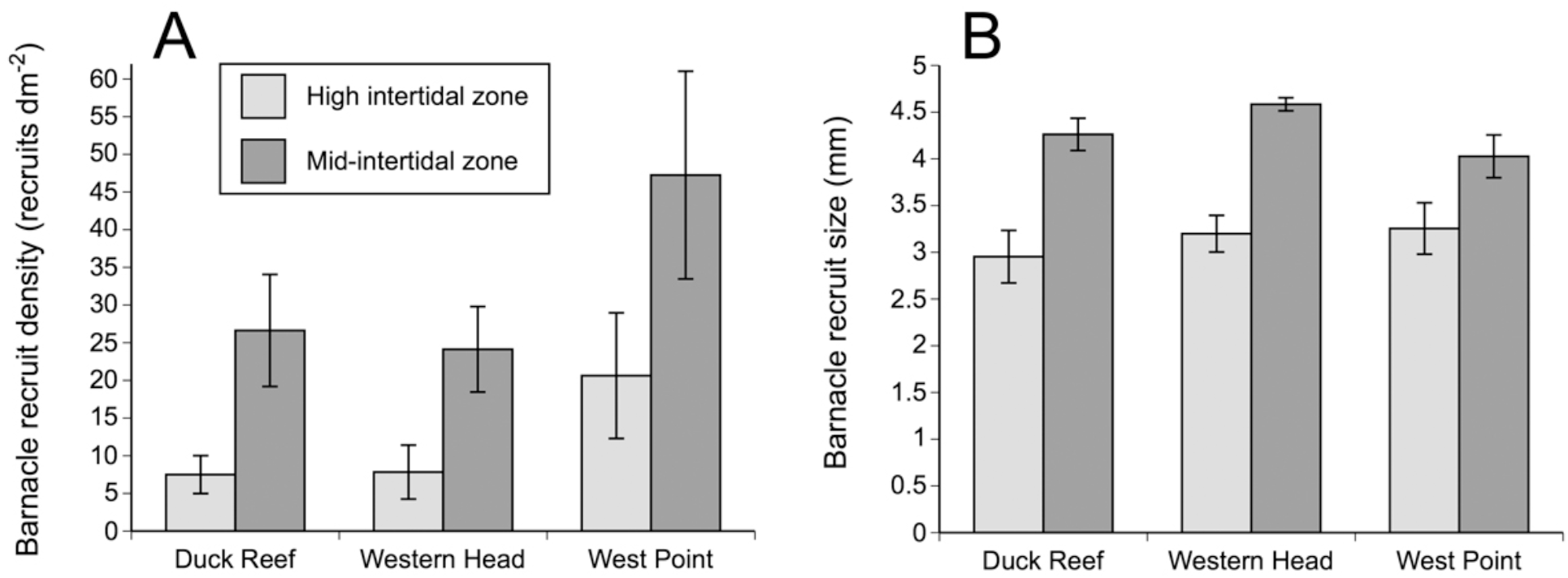
(A) Density and (B) size of barnacle recruits (mean ± SE) at the end of the 2019 recruitment season at the three studied locations on the Atlantic coast of Nova Scotia, Canada.

In early July, barnacle recruit size was significantly higher at the mid-intertidal zone than at the high intertidal zone (*F*_1, 2_ = 39.65, *P* = 0.024), 37 % higher on average (Fig. 2). This outcome was consistent across locations, because the interaction term between elevation zone and location was not significant (*F*_2, 39_ = 1.14, *P* = 0.330). Recruit size was statistically similar among locations (*F*_2, 39_ = 0.88, *P* = 0.424). For recruit size, the denominator degrees of freedom for location and for the interaction term was 39 (instead of 40, as for recruit density) because a quadrat at the high intertidal zone did not have any recruits, so no size value was available for it (that quadrat was included in the analysis of barnacle recruit density because it had a valid value of density: 0 recruits dm^−2^). Thus, the analysis of variance for recruit density included data for 46 quadrats, while the analysis for recruit size included data for 45 quadrats. Visual examples of typical differences in recruit density and size between high and middle intertidal elevations are shown in Fig. 3.

**Figure 3.**
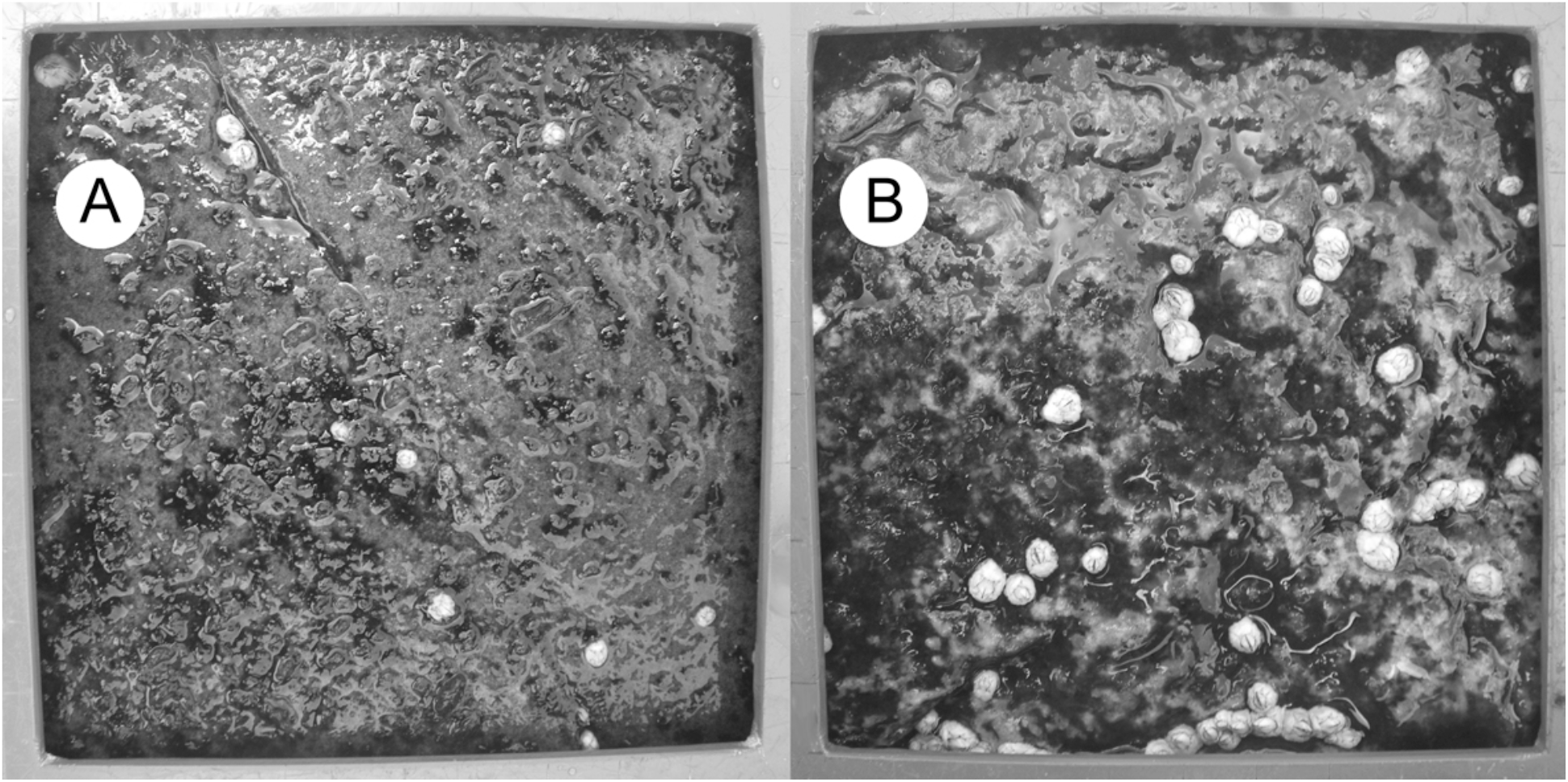
Typical differences in barnacle recruit density and size between (A) high and (B) middle intertidal elevations. The frame bordering each photo belongs to the sampling quadrat. Each side of the quadrat measures 10 cm.

## Discussion

The analysis of the 2019 recruitment season for the barnacle *Semibalanus balanoides* revealed that the density and size of recruits were higher at middle than at high elevations in wave-exposed intertidal habitats on the Atlantic coast of Nova Scotia. These findings agree with the study’s hypotheses, which stemmed from knowledge on basic environmental differences between high and middle elevations. Because of tide dynamics, high elevations often experience more extreme levels of temperature and biological desiccation during low tides than middle elevations (Eckersley and Scrosati 2012, Watt and Scrosati 2013). During low tides, such extremes could have killed a number of recently settled cyprid larvae before they had the opportunity to metamorphose to recruits, in that way limiting recruit density (Shanks 2009). Recruit density may also have been higher at middle elevations because of the longer time such elevations spend underwater, which provides additional time for more pelagic larvae to settle (Minchinton and Scheibling 1993). Abiotic extremes at high elevations could also have limited the growth of established recruits (Bertness 1989, Lathlean et al. 2013), leading to smaller recruits than at middle elevations. As recruits started to appear on the studied coast at similar times at high and middle elevations, the observed size differences between both zones likely reflect differences in growth rates. Lastly, the longer time middle elevations spend underwater also results in a longer period of pelagic food availabilty for recruits, which likely also contributed to the higher recruit sizes at middle elevations.

Despite barnacle recruitment being, on average, nearly 200 % higher at middle than at high elevations in Nova Scotia, values are still lower than those reported for other wave-exposed coasts. For example, the density of *Semibalanus balanoides* recruits at the end of its recruitment season on the NE Atlantic coast reached mean values of 1600–2000 recruits dm^−2^ at some locations in Ireland (Jenkins et al. 2000). On the NE Pacific coast in Oregon and California, recruits of *Balanus glandula* and *Chthamalus dalli* appear throughout most of the year and can reach combined mean densities of 1800 recruits dm^−2^ in just one month (Navarrete et al. 2008). In contrast, for the locations studied in 2019 on the NW Atlantic coast in Nova Scotia, mean density was only 33 recruits dm^−2^ at middle elevations (Fig. 2). For the same three locations (Duck Reef, Western Head, and West Point), mean recruit density in 2014 was 67 recruits dm^−2^ at high elevations (with a highest locationwise density of 108 recruits dm^−2^ at West Point; Scrosati and Ellrich 2018). Assuming for 2014 a 200 % increase in recruit density at middle elevations (as found for 2019), values would still be considerably lower than the highest values found for the NE Atlantic and NE Pacific coasts. Even the highest locationwise value of recruit density found at high elevations in 2014 (212 recruits dm^−2^ at a wave-exposed location north of those studied in 2019; Scrosati and Ellrich 2018) would yield a lower value than for the NE Atlantic and NE Pacific coasts even if increased by 200 %. Farther south on the NW Atlantic coast (in Maine, USA), the mean recruitment of *S. balanoides* in wave-exposed locations between 1972-1974 ranged between 800–1100 recruits dm^−2^ (Menge 1978). Thus, wave-exposed intertidal environments on the NW Atlantic coast in Nova Scotia show relatively moderate values of barnacle recruitment.

When measured at the same relative elevation (relative to the full intertidal range), differences in barnacle recruitment between coasts are determined by a variety of factors. Among the seemingly most important ones, studies have identified surf zone hydrodynamics (Shanks et al. 2017, Shanks and Morgan 2019), coastal upwelling (Menge and Menge 2013, 2019), seawater temperature and pelagic food abundance (Cole et al. 2011, Scrosati and Ellrich 2018), and coastal winds affecting larval transport (Bertness et al. 1996), for example. Therefore, a multivariate study focusing simultaneously on these factors should help to unravel what drives the notable differences in barnacle recruitment observed among the wave-exposed shores so far studied. Ultimately, the observed differences in the supply of recruits suggest that the intensity of interspecific interactions between barnacles and other intertidal species may differ considerably among these coasts.

## Acknowledgements

I am grateful to Matthew J. Freeman for his help with the experimental setup and to the Natural Sciences and Engineering Research Council of Canada (NSERC) for funding this project through a Discovery Grant.

